# Performance comparison of probe-capture enrichment sequencing of viruses in wastewater influent and settled solids from California’s Central Valley

**DOI:** 10.64898/2026.10.02.756336

**Authors:** Amit Lavon, Julian M. Pender, Rachel Olson, Titash Chatterjee, Colleen C. Naughton, Priyanka Prashar, Scott Kuersten, Jonathan A. Eisen, Katrine Whiteson, Heather N. Bischel

**Affiliations:** Dept of Molecular Biology and Biochemistry, University of California, Irvine; Biochemistry, Molecular, Cellular, and Developmental Biology (BMCDB) Graduate Group, University of California, Davis; Dept of Civil and Environmental Engineering, University of California, Davis; Dept of Civil and Environmental Engineering, University of California, Merced; Illumina, Inc., San Diego, CA, USA; Genome Center, Dept of Evolution and Ecology, and Dept of Medical Microbiology and Immunology, University of California, Davis

## Abstract

Wastewater sequencing using probe-capture enrichment of public health-relevant viruses is a powerful, non-intrusive approach to monitor infectious diseases, but performance depends on sample type and processing approach. We compared two processing pipelines from wastewater influent samples collected at three timepoints across three wastewater treatment facilities. Each sample was split for parallel processing, with viruses concentrated (1) directly from whole influent using nano-sized affinity particles and (2) by settling and dewatering the solids fraction. Combined DNA/RNA extracts were prepared into libraries using a pre-commercial Illumina protocol, then probe-capture enrichment was performed with Illumina’s Viral Surveillance Panel (VSP) v1 (66 targets) and v2 (242 targets). Sequencing depth was increased for VSP v2 to accommodate the expanded target set. On average, 160 ± 146 viral species were detected per sample, with wastewater fraction (sample type) explaining 19% of the variance in viral abundances. Solids contained a higher abundance of DNA viruses compared to RNA viruses, while influent exhibited the inverse. Metagenomic analysis of the VSP v2 sequencing data detected more viral species in influent (median 358) than in solids (median 184), but probe-capture enrichment produced more comparable target-virus readouts across fractions, supporting the use of either fraction for wastewater-based epidemiology.

## Introduction

Wastewater sequencing has emerged as a useful tool for public health surveillance. It enables tracking disease outbreaks, monitoring infection trends, and identifying emerging pathogens. Different sequencing methods result in different pathogen detection rates. Tiled amplicon wastewater sequencing has received attention for its applications in COVID-19 monitoring (Tiwari et al. 2023; Karthikeyan et al. 2022). Our group (Rothman et al. 2021 and 2023) and others (Child et al. 2023) have demonstrated the advantages of using probe-capture enrichment sequencing over untargeted shotgun sequencing, finding significant improvements in the percentage of viral reads, genome coverage, and mean coverage depth. Enrichment using tailored hybridization probes increases the relative read abundances selectively, increasing the percentage of specific pathogen reads by orders of magnitude (Kantor and Jiang 2024).

The choice of sample type, virus concentration methods, and extraction methods influences sequencing outcomes. High sensitivity of SARS-CoV-2 detection by PCR has been achieved through processing of wastewater influent and settled solids, with primary settled solids containing approximately three orders of magnitude higher SARS-CoV-2 RNA concentrations than influent on a mass-equivalent basis, making solids an attractive matrix for sensitive ddPCR-based monitoring (Kim et al. 2022). In the context of metagenomic sequencing, however, the balance between viral enrichment and background nucleic acids becomes crucial, and several groups have used hybrid-capture panels to examine how sample type and processing impact viral read fractions. In a study evaluating different wastewater sample processing methods in California, Jiang et al. (2024) found that nucleic acid extraction from solids pelleted out of whole influent yielded a lower percentage of viral reads (0.17%) than any of the three influent extraction methods, and the highest virus relative abundance was achieved using an InnovaPrep ultrafiltration pipette (1.82%), with *Astrovirus* and *Polyomaviruses* as the most abundant human viruses across the tested protocols. In a larger-scale hybrid capture study, Wolfe et al. (2026) applied a broad viral panel to wastewater liquids and found that influent-derived samples consistently exhibited higher viral read fractions than sludge supernatant, with several influent sites exceeding 25% viral reads. Further evaluation of sample processing methods will provide value in understanding associated tradeoffs when characterizing the pathogen profiles of wastewater.

Beyond methodological comparisons, several studies have utilized hybridization probe-capture approaches to survey viral diversity in wastewater. A survey of southern California wastewater using the Illumina Respiratory Virus Oligo panel enabled enhanced detection of coronaviruses, influenza viruses, adenoviruses, and bocaviruses compared to unenriched metagenomic sequencing of the same samples (Rothman et al. 2021). Detection of variants required tiled amplicon sequencing and enabled detection of several SARS-CoV-2 sublineages in wastewater before clinical detection (Rothman et al. 2024; Karthikeyan et al. 2022). McCall et al. (2023) used the ViroCap probe panel in Houston, Texas, finding greater congruence in viral composition between wastewater treatment plant samples than between building-level samples. Their analysis revealed *Astroviridae* and *Adenoviridae* as the most abundant viral families in the treatment plant samples. Similarly, a broader survey across Houston and El Paso, employing the TWIST Comprehensive Virus Research Panel, consistently detected strains belonging to *Astroviridae*, *Adenoviridae*, and *Polyomaviridae*, alongside public health-relevant viruses like SARS-CoV-2, influenza, and monkeypox (Tisza et al. 2023).

The application of enrichment sequencing with probe-capture sets has also enabled the identification of specific viral strains. In a study of California’s Bay Area wastewater, Crits-Christoph et al. (2021) successfully detected SARS-CoV-2 alleles that correlated with clinical data. However, Rothman et al. (2021) and Rothman and Whiteson (2022) showed that even when probe-capture is used to enrich for respiratory viruses, wastewater libraries were still dominated by off-target plant-infecting viruses, particularly tobamoviruses.

In this study, we experimented with Illumina’s Viral Surveillance Panel (VSP), versions 1 and 2, to compare their pathogen detection potential. VSP is a probe enrichment panel designed to capture suites of viruses of significance to public health. Version 1 (v1) of the panel is designed to capture 66 viral genomes, while version 2 (v2) is designed to capture 242 viral genomes including all those in v1. We set out to explore the compatibility of each panel with both wastewater influent and settled solids. Three locations were each sampled at three timepoints, resulting in nine wastewater samples. Each sample was processed in two ways, by using nano-sized microbiome affinity particles for virus concentration (referred to herein as *influent* samples) and by collecting settled solids for direct extraction (referred to herein as *solids* samples). Sequencing of enriched libraries was conducted in two batches, one using VSP v1 with ∼6.7 million read pairs expected per sample and one using VSP v2 with ∼67 million read pairs expected per sample to support its larger target enrichment set. Throughout this manuscript, the term sequencing configuration refers to the specific combination of VSP panel version and sequencing depth used for a given analysis.

## Results

### Study Samples

We compared four sequencing configurations, [solids, influent] x [VSP v1, VSP v2], with 9 samples in each group. VSP v1 samples averaged 8.6 ± 4.6 million (M) read pairs pre-QC and 8.1 ± 4.6M read pairs post-QC. VSP v2 samples averaged 76 ± 14M read pairs pre-QC and 75 ± 14M read pairs post-QC (Figure 1a). Considering sample type within each batch, the median sequencing depths for v1 were 4.8M for solids and 10M for influent, while for v2 they were 64M for solids and 85M for influent. The proportion of reads assigned to viruses ranged from 0.05% to 50% when including sequences associated with positive control spikes: bovine coronavirus (BCoV) was added before extracting nucleic acids whereas RNA from Influenza A and DNA from Adenovirus Type 1 were added to the DNA & RNA extract for each sample (Table S1 and Table S2). The proportion of reads assigned to viruses ranged from 0.014% to 0.55% when excluding sequences associated with the spiked positive controls (Figure 1b). The average viral read ratios without positive control spikes were 0.045% of the reads per sample for Solid VSP v1, 0.14% for Solid VSP v2, 0.32% for Influent VSP v1, and 0.18% for Influent VSP v2.

**Figure 1:**
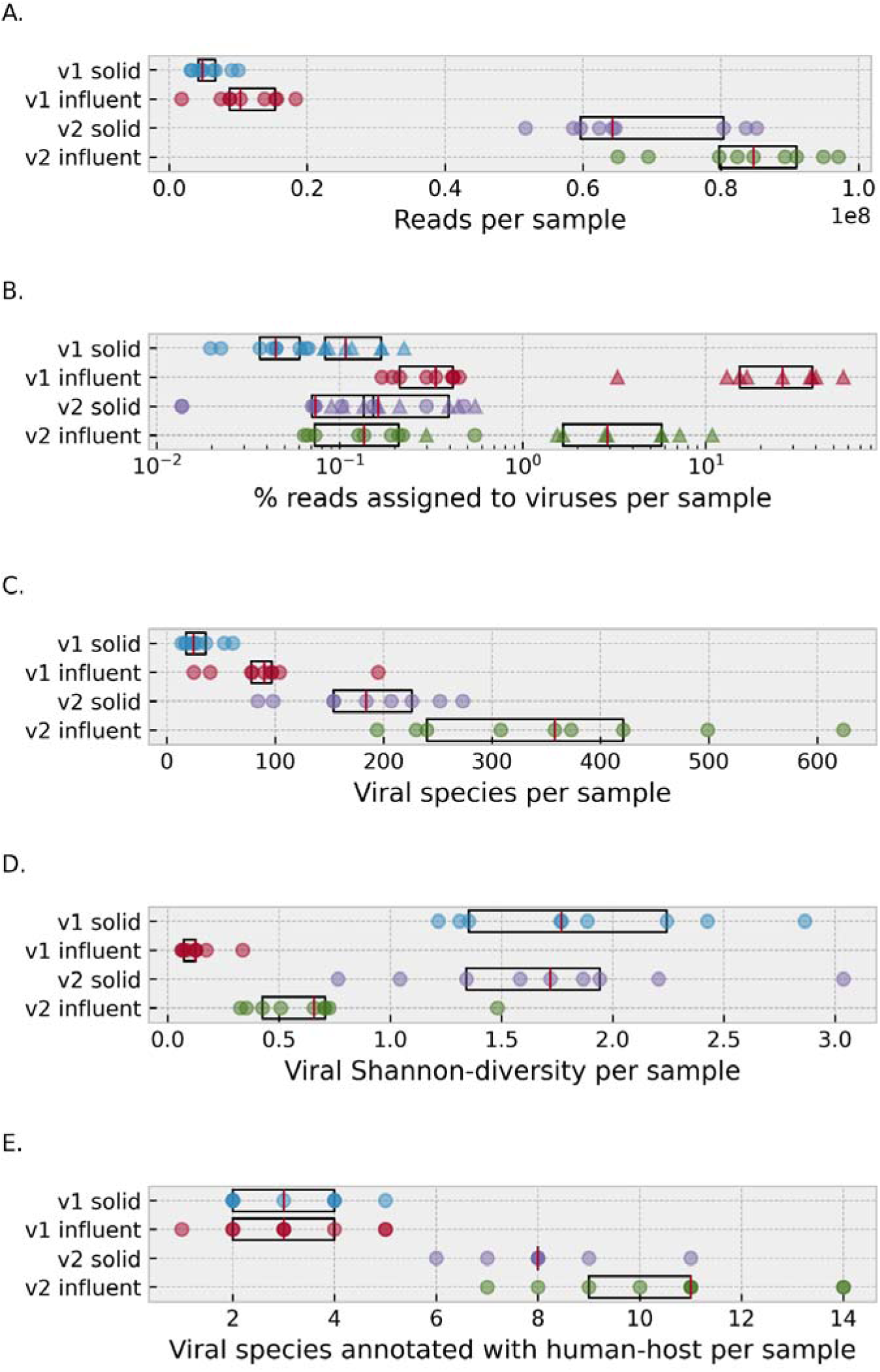
Sample characteristics and read distributions grouped by sequencing configuration (v1/v2) and sample type (solid/influent). (A) Distributions of total number of reads post-QC, with data including sequences associated with the positive control spikes. (B) Percentage of reads assigned to viruses per sample, where triangles indicate data with positive control sequences retained, while circles indicate data with these sequences removed. Note the x-axis is log scale. (C) Viral species per sample as determined by Kraken. (D) Shannon-diversity of viral species. (E) Number of viral species that were annotated with human-host in RefSeq’s metadata. Data in panels C–E represent results excluding sequences associated with the positive control spikes.

On average, we detected 160 ± 146 viral species per sample. The median number of viral species identified per sample by Kraken was 25, 90, 184, and 358 for v1 solid, v1 influent, v2 solid, and v2 influent, respectively (Figure 1c). The Shannon diversity of viral relative abundances ranged from 1.3 to 4.5 bits (Figure 1d). The number of viral species with a human host annotation using available RefSeq metadata ranged from one to 14 (Figure 1e). Of the viruses detected that we associated with the VSP v2 panel targets using a graph data structure approach, 36 viral species were detected at least once in the solids and 46 viral species were detected at least once in the influent (Table S3). The VSP v1 pipeline enabled detection of 23 of the panel targets in the solids and 26 in the influent. In a rarefaction analysis, the number of viral species in the v2 samples was similar to the v1 samples in the v1 read count area (Figure 2). At higher read counts obtained via the VSP v2 pipeline, rarefaction curves generally plateaued at somewhat higher numbers of viral species in influent compared to solids, though not consistently. The number of viral species identified generally increased with the number of reads, with modestly higher species per read identified in influent relative to solids (Figure S1).

**Figure 2:**
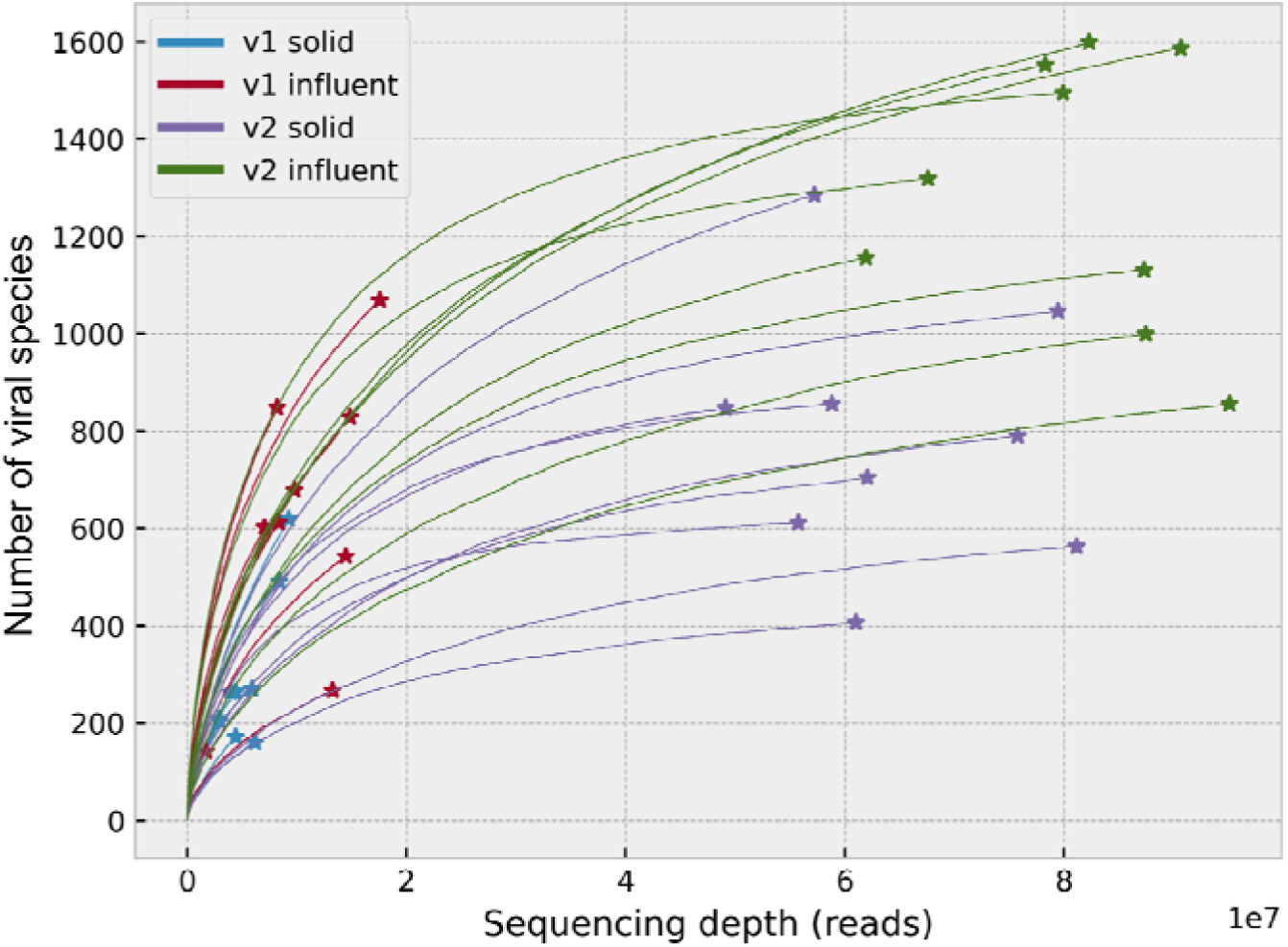
Rarefaction curves of the 36 samples, using raw read assignments and including sequences associated with positive control spikes.

### Viral relative abundance comparison

We estimated microbial relative abundances using Kraken (see methods) and focused on the groups under *“Viruses”*. We recovered positive control spikes (*Betacoronavirus 1, Human mastadenovirus C*, and *Alphainfluenzavirus influenzae*) in most samples (Table S2). Excluding sequences associated with positive control spikes, the most abundant viral species on average were *Human astrovirus*, *Carjivirus communis* and *Betapolyomavirus secuhominis* (Figure 3). For the VSP v1 samples, the dominant viral species in solids were Human astrovirus and *Carjivirus communis*, while the dominant species in influent were Rabbit coronavirus HKU14 and Human astrovirus. Rabbit coronavirus HKU14 (RbCoV HKU14) is a genetically distinct rabbit coronavirus that belongs to the same subgenus, *Embecovirus*, as bovine coronavirus. Its apparent detection is therefore consistent with, but does not establish, cross-classification of reads originating from the bovine coronavirus positive-control spike-in. We found no published reports or publicly-available sequence or RT-PCR-confirmed detections of RbCoV HKU14 in the United States. In the VSP v2 samples the dominant viral species in solids were Human mastadenovirus F and *Jouyvirus ev017* (initially known as crAss-like phage), while the dominant species in influent were Human mastadenovirus F and *Dependoparvovirus primate1* (formerly known as Adeno-associated virus). The Pearson correlation coefficients for viral species relative abundance within each sequencing pipeline indicate greater similarity between solids and influent of results for the VSP v1 pipeline compared to VSP v2 (Figure 4). When considering the 12 most relatively abundant viral species across all sample groups, the relative abundance of some viruses correlated with VSP sequencing pipeline, such as *Dependoparovirus*, and some viruses correlated with sample type, such as tomato brown rugose fruit virus (Figure 5).

**Figure 3:**
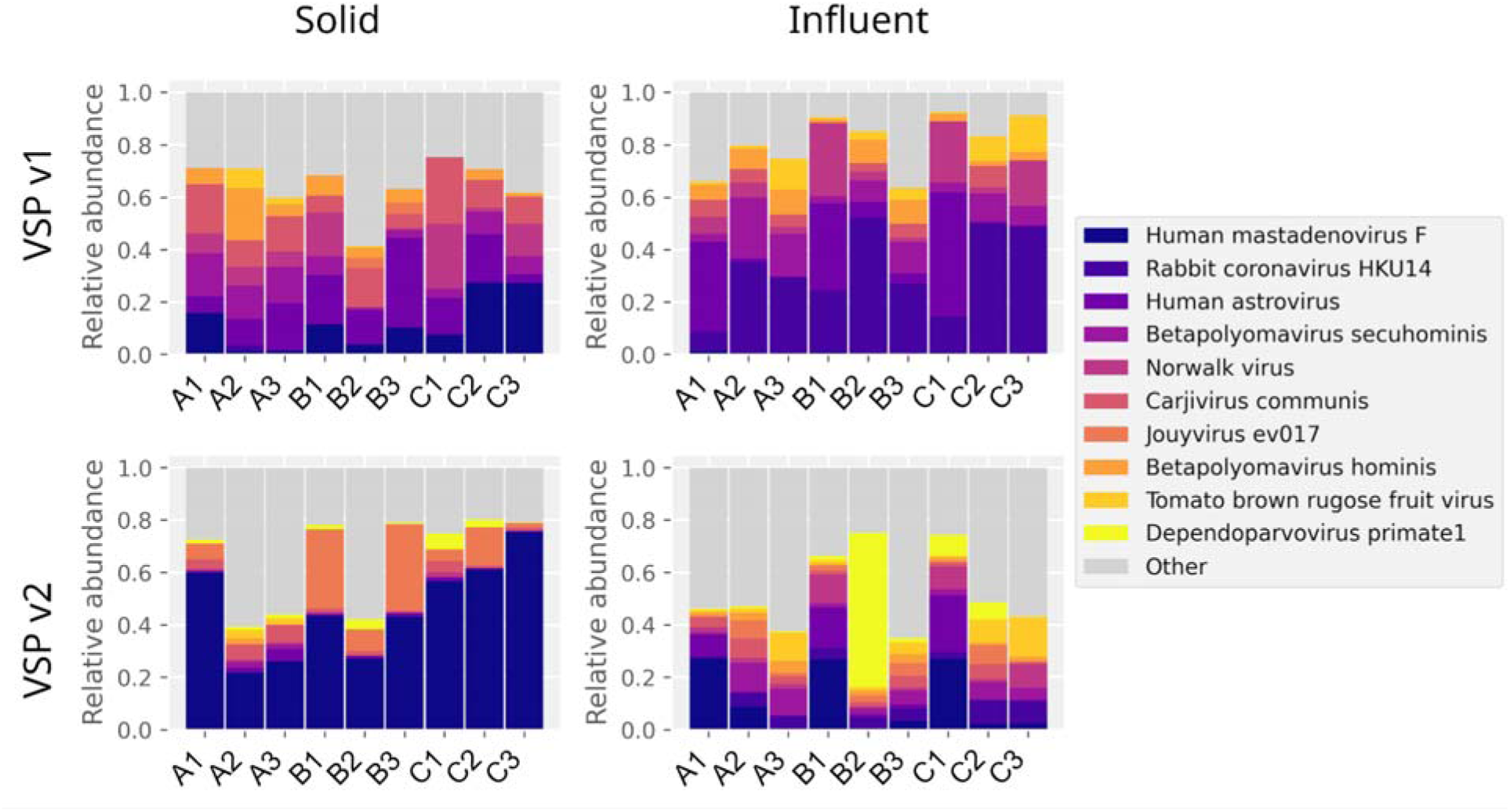
Viral relative abundances per sample, without sequences associated with positive control spikes. Samples A1-A3, B1-B3, and C1-C3 represent three sequential collections from Sites A, B, and C, respectively; numbers indicate collection order. The 10 most abundant species across all samples are colored individually, and the rest are grouped as “Other”.

**Figure 4:**
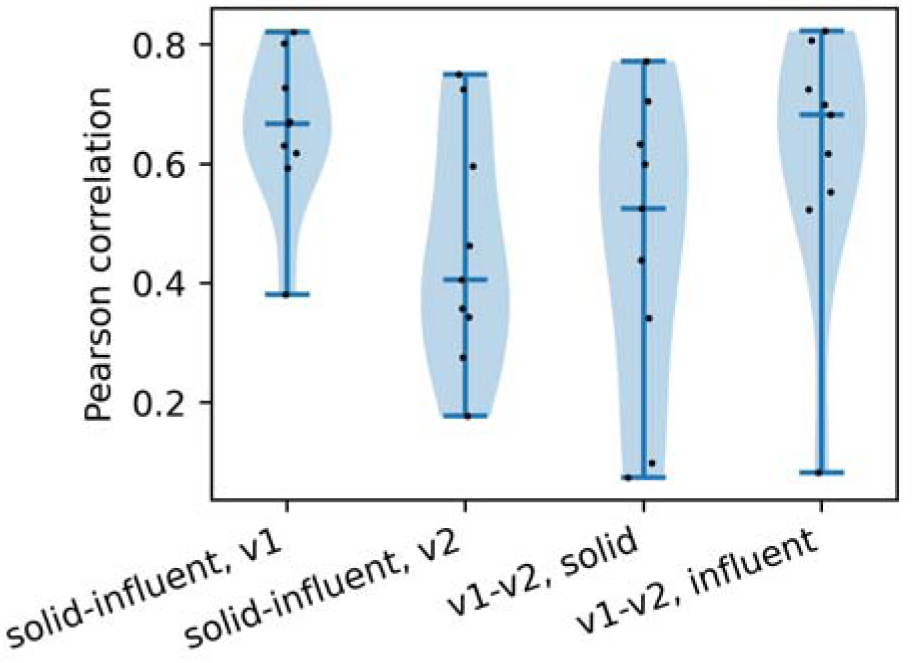
Viral relative abundance correlations across sample types and sequencing methods. Each point represents a single sample, comparing two sample types (left two correlations) or two sequencing methods (right two correlations).

**Figure 5:**
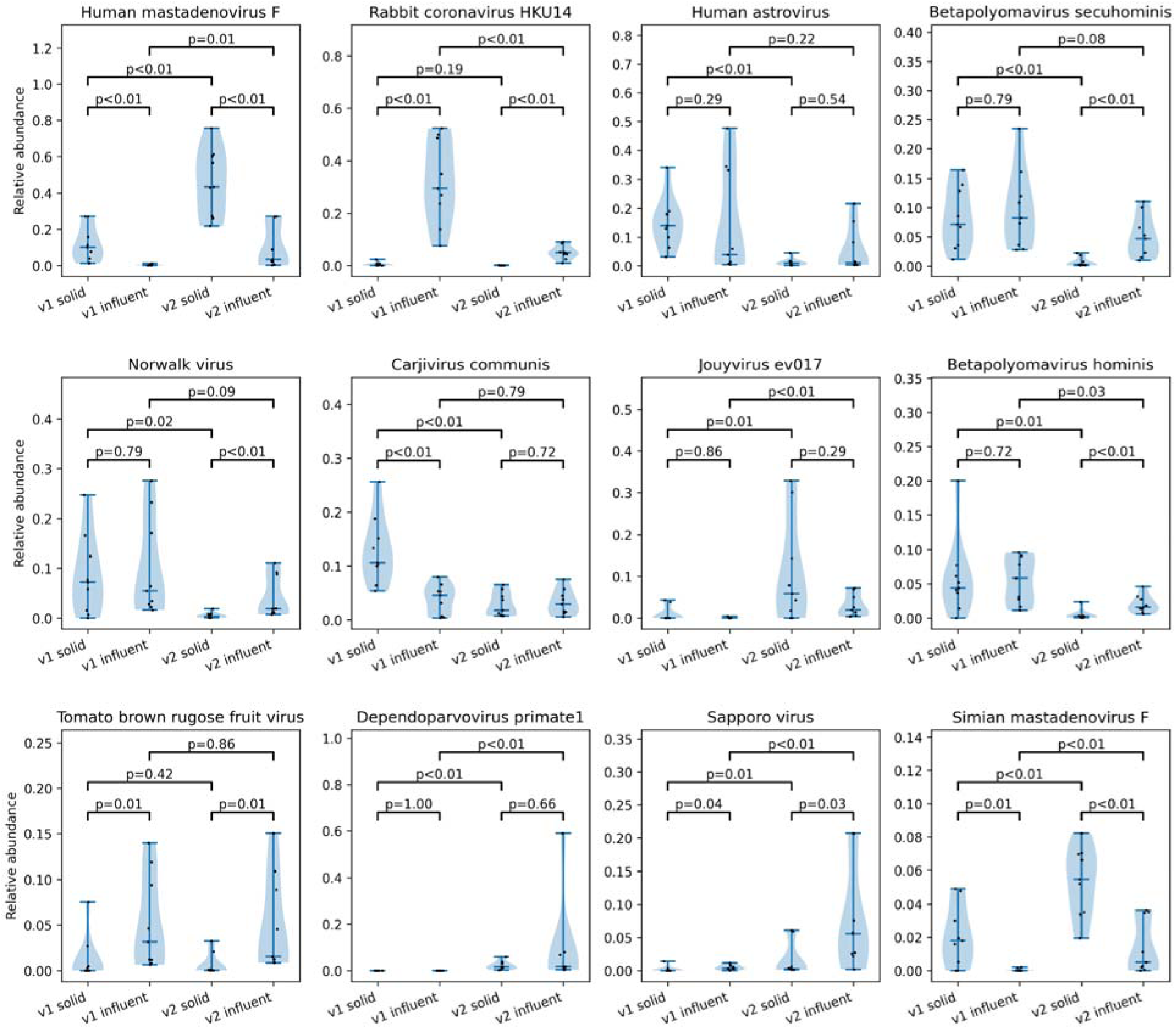
Comparison of relative abundances of the 12 most abundant species across sample groups. Human mastadenovirus F, Human astrovirus, Betapolyomavirus secuhominis, Norwalk virus, Betapolyomavirus hominis, and Sapporo virus are associated with the VSP panel enrichment targets. The species with the highest mean relative abundances are shown, broken down by sample group. Mann-Whitney U tests were performed between pairs of groups, and the P-values are shown.

To assess the influence of sequencing configuration, sample type, and sample location on relative abundances of viruses detected, we conducted a multivariate analysis (Figure 6). Using a PERMANOVA test on the viral relative abundances without spike, we found that the sequencing configuration accounted for 23% of the variance (p=1×10^-6^), sample type (solid/influent) accounted for 19% (p=1×10^-5^), and location accounted for 8% (p=0.12). For comparison, we also applied PERMANOVA on the relative abundances of all domains, and found the sequencing configuration accounted for 8% of the variance (p=0.028), sample type (solid/influent) accounted for 37% of the variance (p=1×10^-6^), and location accounted for 17% of the variance (p=0.0031). Looking at the top-level distributions, the samples were dominated by bacteria, unknowns, and the *Betacoronavirus 1* positive control (Figure S2). Bacterial relative abundances were less affected by the sequencing configuration than by the sample type (Figure S3 and S4).

**Figure 6:**
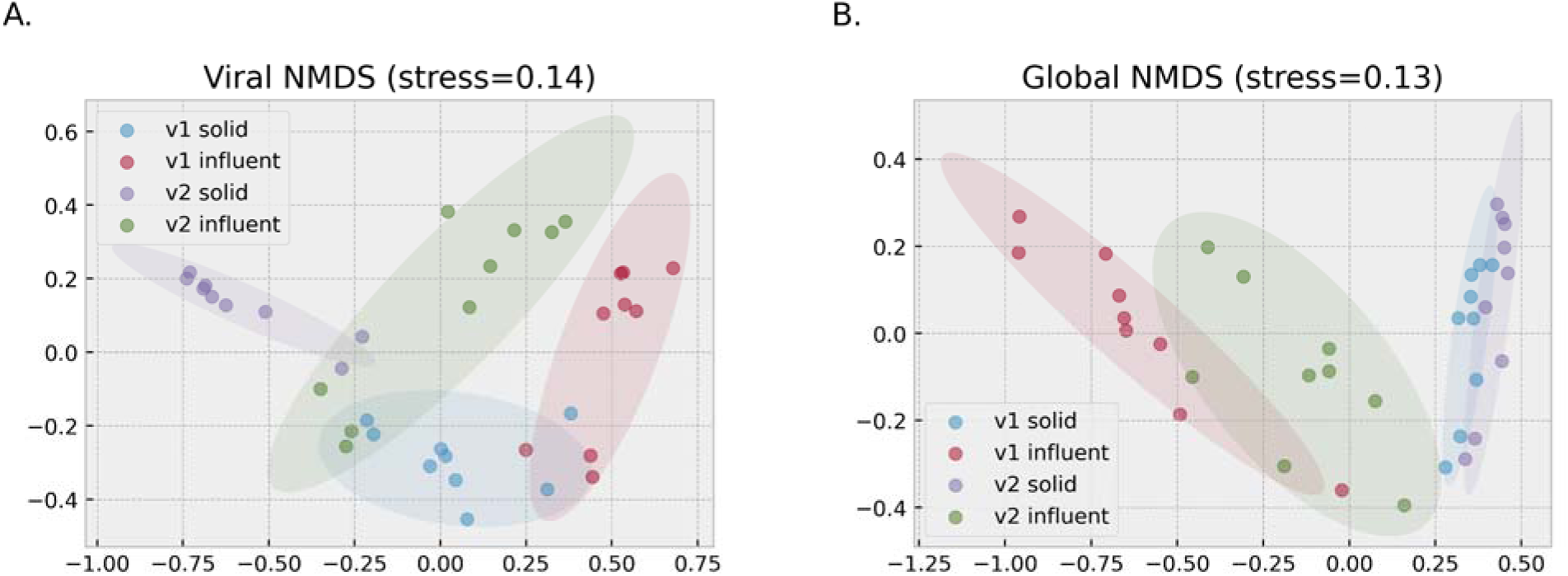
Non-metric multidimensional scaling (NMDS) of (A) the viral component of the sequence data, and (B) all the species of the sequence data. The dissimilarity score used for the NMDS is pairwise Bray-Curtis on the relative abundances, without assumed spikes. Samples are colored by group.

### Genome coverage

To validate the identification of spike-in controls and selected abundant *in situ* viruses, we evaluated how genomes for the viruses were covered by the samples’ reads. We mapped each sample’s reads, including sequences associated with positive control spikes, to a representative RefSeq reference genome for the virus of interest. We then collected the reads that mapped uniquely and plotted the coverage by sample type (Figure 7). The total non-uniquely mapped reads comprised less than 0.01% of the reads. Consistent with expectation, lower coverage depth was evident for results from the v1 sequencing pipeline compared to v2. Coverage depth of the post-extraction positive control spike of Adenovirus Type 1 (using Human mastadenovirus C as a representative genome) was similar in pattern and order of magnitude between the influent and solids samples, within each sequencing pipeline (Figure 7B). Coverage depth of the pre-extraction positive control spike Bovine coronavirus was substantially lower for solids samples compared to influent samples (Figure 7C and 7D), reflecting differences in concentration of the virus spiked in during sample preparation, recovery efficiencies, and sequencing depth obtained for each sample type. For the *in situ* viruses that were evaluated, the solids samples generally exhibited lower average coverage depths compared to the influent samples, with the exception of Human mastadenovirus F (Figure 7E through 7J).

**Figure 7:**
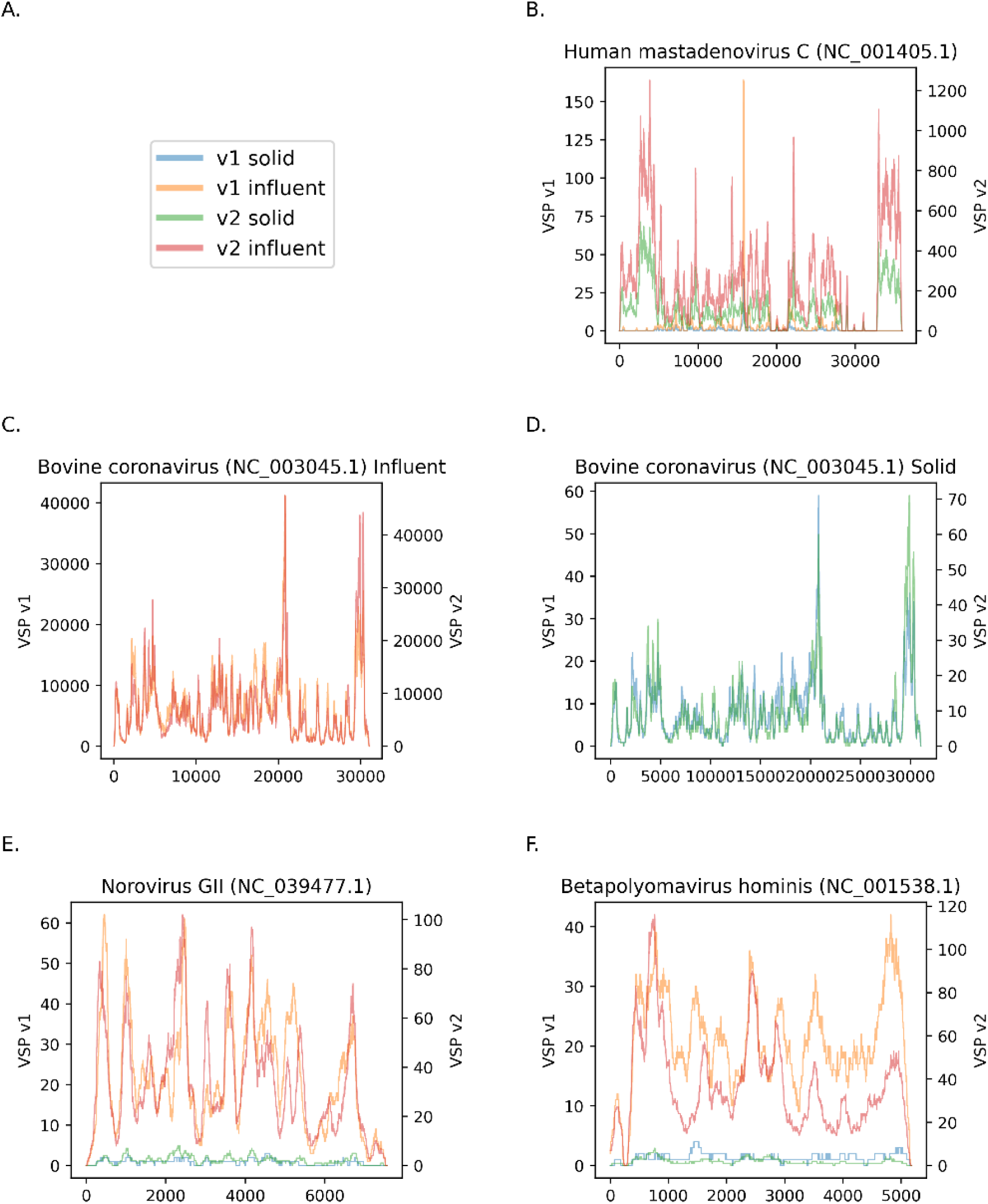
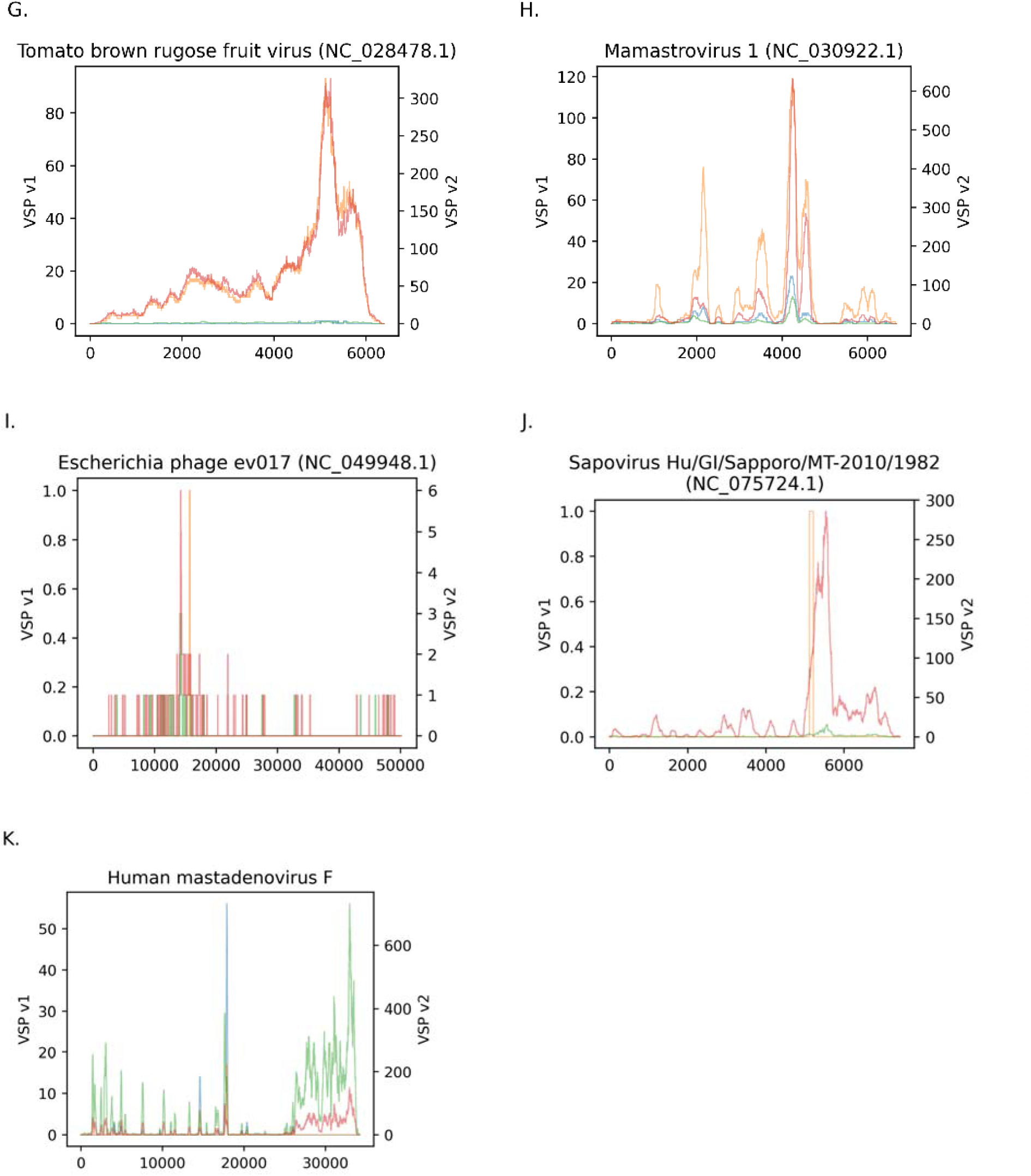
Genome coverage per viral species, where the x-axis is the location along the reference genome, and the y-axis is average reads per base. (A) Uniquely mapped reads were aggregated by sequencing configuration and sample type (v1 solid in blue, v1 influent in orange, v2 solid in green, and v2 influent in red). (B) Human mastadenovirus C was selected to represent the post-sample processing control spike of Adenovirus Type 1. (C-D) Bovine coronavirus was a pre-sample processing control spike in all sample groups. (E-J) Virus names are different from those in Figure 3 because they are from Refseq, not Kraken.

The evaluation of genome coverage for select viruses was also intended to assess whether identification may have resulted from incidental similarity between portions of viral genomes and other sequences. One case that requires further evaluation is *Escherichia* phage EV017 (Jouyvirus), which was the 5th most relatively abundant species on average. Part of the highly abundant intestinal and environmental phages initially referred to as crAss-like (Smith et al. 2023), Jouyvirus are commonly co-detected with *Carjivirus communis* (formerly referred to as the prototypical crAssphage). We analyzed the reads that were classified as Jouyvirus by Kraken using BLAST. Out of the 2008 reads, 1045 (52%) had hits of any type of Jouyvirus. Out of those, 1023 had hits from EV017. However, average coverage of EV017 was as low as 1 read per base, with most of the genome not being covered at all (Figure 7). The other most common hits were *Escherichia* phage ESSI2_ev239 and *Escherichia* phage vB_EcoM_G4507. Running the coverage analysis on those two resulted in no coverage at all.

Coverage analysis of the two most common bacterial species gave a relatively uniform coverage depth, 1,000 for the v1 sequencing configuration and 10,000 for v2 (Figure S5).

### Comparison of RNA and DNA content between sample types

To compare the relative proportion of DNA and RNA viruses identified in influent and solids within each sequencing approach, we reviewed the nucleic acid genome-type of each phylum identified (Table S4) and summed the relative abundances by that annotation (Figure 8). In both VSP v1 and v2, the solids samples contained higher relative abundances of DNA viruses and lower relative abundances of RNA viruses on average (Figure 8). This pattern was also observed when considering only viral species enriched by the VSP v2 and detected at least once (Table S5): among 27 RNA viruses, 13 were detected in solids and 22 in influent, while among 24 DNA viruses, 15 were detected in solids and 19 in influent (Table S3). Additional samples are needed for meaningful statistical evaluation of these observations.

**Figure 8:**
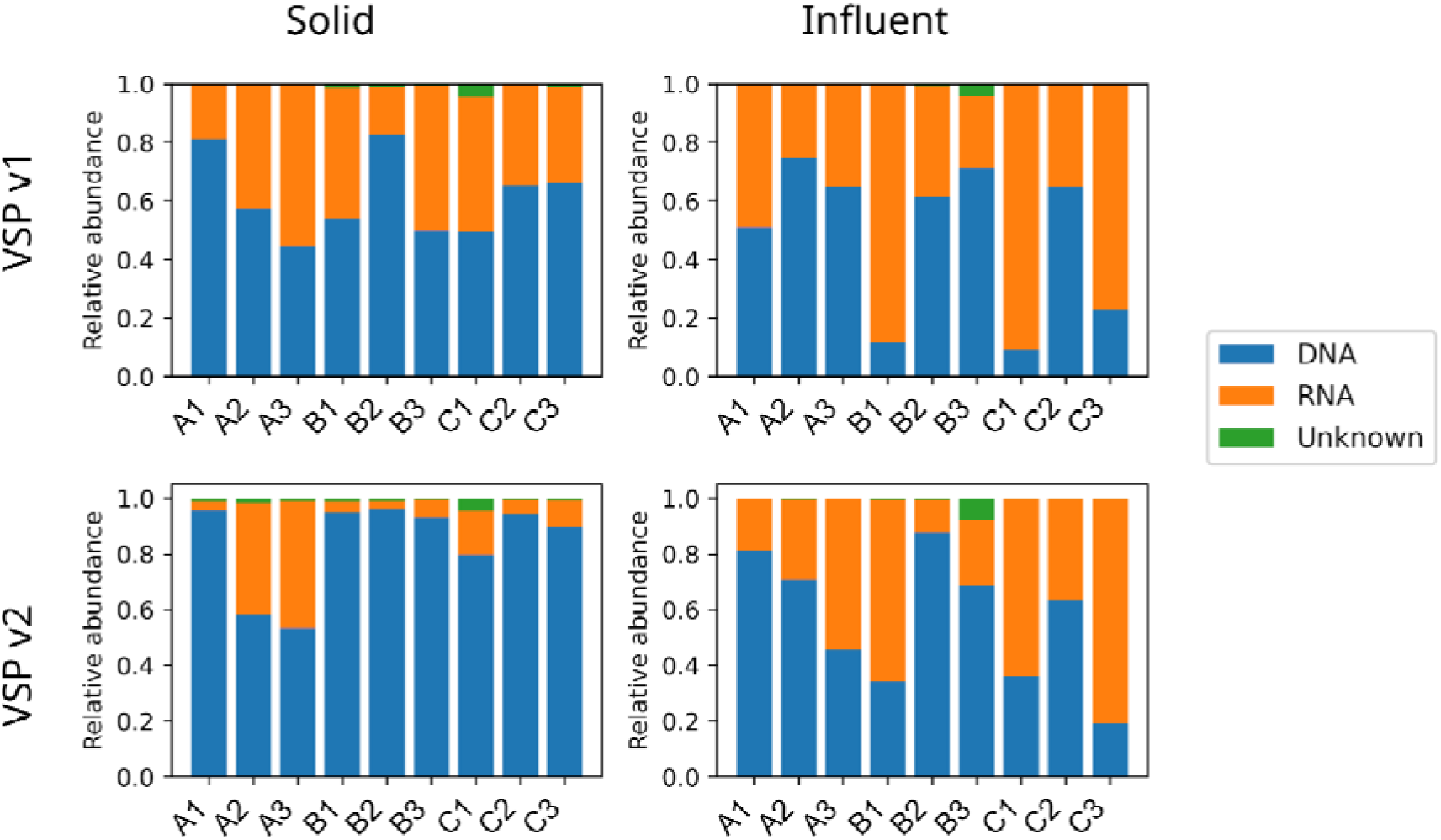
Relative abundances of viruses with RNA or DNA genomes, based on phylum assignment, excluding sequences associated with positive-control spikes. Samples A1-A3, B1-B3, and C1-C3 represent three sequential collections from anonymized Sites A, B, and C, respectively; numbers indicate collection order. Taxa with unknown or unannotated phyla are grouped as “Unknown”. Only the 0.24% of reads mapping to viruses by Kraken are included. Of the 13 viral phyla listed in Table S4, eight are DNA viruses and five are RNA viruses.

To extend the analysis to non-viral RNA, we mapped the reads of each sample to the GreenGenes dataset of bacterial and archaeal rRNA sequences (DeSantis et al. 2006). The percentage of reads mapped successfully to GreenGenes ranged from 6% to 36% (Figure 9). In both v1 and v2, the median ratio in solids was lower than the median ratio in influent. We compared the ratio distributions of solid and influent using Mann-Whitney test, yielding p=0.66 for v1 and p=0.04 for v2.

**Figure 9:**
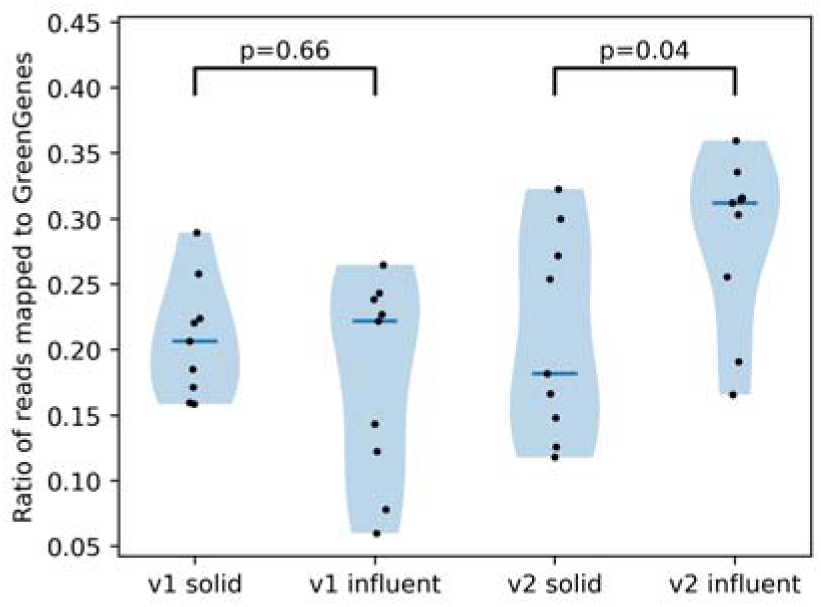
Distributions of ratio of reads mapped to GreenGenes. p-values of Mann-Whitney tests compare solids and influent within each VSP version.

## Discussion

In this effort, we explored the virome of wastewater samples from California’s Central Valley region, using two different sample processing methods for each of two sequencing configurations. We observed that the sequencing configuration had a larger impact on viral relative abundance (23% of variance) than the sample processing method (19% of variance, Figure 6). Because the VSP v2 submission was sequenced at a greater depth than the VSP v1 submission and v2 samples resulted in approximately nine times more reads on average, these effects cannot be directly attributed to the broadened panel. Rarefaction analysis showed that at the maximum read depth of VSP v1, the number of viral species identified differed little between solids and influent (Figure 2). In contrast, at the higher read depths of VSP v2, rarefaction curves often plateaued earlier for solids, indicating fewer viral species than in influent.

In terms of relative abundance, over 95% of each sample’s reads were assigned as bacteria, spike, or unclassified (Figure S2). The low proportion of reads assigned to viruses (0.014% to 0.55%) is within range of previous findings (Rothman et al. 2021, Rothman et al. 2024, Jiang et al. 2024 and reviewed by Kantor and Jiang 2024). The most abundant viral species in solids were Human mastadenovirus F, Human astrovirus, *Carjivirus communis* and *Jouyvirus ev017*. In influent the most common species were Rabbit coronavirus HKU14, Human astrovirus, *Betapolyomavirus secuhominis* and Norwalk virus. Genome coverage analysis of the most abundant viral species showed similar distributions across the different sample types. These taxa, which include both human enteric viruses and bacteriophages, are commonly detected in surveys of the wastewater virome (McCall et al. 2023; Tisza et al. 2023; Jiang et al. 2024). The dominant species herein differed from wastewater influent sequencing in Rothman et al. 2021 and Rothman et al. 2024, where tobamoviruses were most abundant in both unenriched shotgun sequencing and enriched with the Illumina Respiratory Virus Oligo Panel.

We focused further analysis on the differences between pre-enrichment sample processing workflows within each enrichment-sequencing configuration. The number of viral species identified in each sample correlated with the number of reads, but was generally higher in the influent samples, in terms of species per read (Figure S1). Although equi-molar library pools were prepared within each sequencing pipeline, the influent samples had 112 ± 122% more reads than their solid counterparts in the VSP v1 pipeline, and 27 ± 28% more reads than matched solids samples in v2. The unequal read depths can bias detections towards the higher depth group, especially for rare detections, with higher noise rates from the computational pipeline at higher read counts also a likely additional contributor. Nevertheless, the median number of viral species detected per sample in influent (358) was nearly twice that in solids (184) by the VSP v2 pipeline, and over three times greater in the VSP v1 pipeline (90 vs. 25). We note that changing Bracken’s minimum read threshold drastically affects the number of species identified. We set the threshold to 10 here, which means that a species must have at least 10 reads assigned to it in order to be reported. While the “best” value for this threshold is not clear, as it has to balance between eliminating noise while retaining the informative hits, a threshold of 10 helps avoid spurious hits.

When we examined the viral relative abundances by their genome type, we found that a broad range of DNA and RNA viruses were detected in both sample types and by both sequencing pipelines (Tables S4 and S5). Solids contained on average a higher relative abundance of DNA viruses than RNA viruses, while influent contained a higher relative abundance of RNA viruses than DNA viruses. Mapping the reads to GreenGenes provided a proxy for rRNA amounts in the samples, which were higher in influent as well, although not significantly.

The effect of the sample processing method on the detection of VSP-enriched viruses may be less than its effect on total viral species. With nearly 50 public health-relevant viral pathogens detected across the samples, 46 were detected in influent and 36 in solids by the VSP v2 pipeline. In other words, VSP enrichment may provide greater consistency across different sample processing methods. Application of virus-enrichment sequencing of wastewater influent and settled solids in longitudinal studies are needed to further evaluate this effect.

Wastewater sequencing with probe-capture enrichment of viruses shows great promise to enable the identification of low relative abundance, public health-relevant viruses. Tailored bioinformatics pipelines for enrichment targets will further enable rapid and accessible communications of wastewater sequencing results to inform public health. Our findings demonstrate that two hybridization probe-capture enrichment sequencing pipelines were compatible with both whole influent and settled solids samples. We demonstrate that the choice of sample processing methods is an important factor to consider when sequencing wastewater, as it can influence the types of viruses recovered.

## Methods

### Sample collection and nucleic acid extraction overview

This study analyzes total nucleic acid extracts from composite wastewater influent samples collected from three municipal wastewater treatment facilities (Sites A-C). Three aligned sampling timepoints in 2022 were selected. The first sampling event occurred in mid-April (April 14 for Site A, April 7 for Site B, and April 15 for Site C), followed by two additional sampling events on November 16 and November 30 for all sites.

Sample extraction methods for both influent and settled solids are described in prior publications (Daza-Torres et al. 2023; Muralidharan et al. 2025 for influent; Kadonsky et al. 2023 for settled solids) and are summarized in the following two sections. As a process control, all samples were spiked with bovine coronavirus (BCoV) prior to concentration and extraction, with spike-in concentrations differing between solids and influent fractions. Sample extracts for both sample types were stored at -80 °C until use for this study. After storage, all extracts were also spiked with whole genome extracts of Influenza A H1 (RNA virus control, expected to have low prevalence during the study period) and Adenovirus Type 1 (DNA virus control). Bioinformatics analyses are presented both with and without the spike data included.

### Influent sample processing

From each ∼1 L wastewater influent sample, a 50-mL aliquot was collected before solids settling and transported on ice to UC Davis. Aliquots were pasteurized at 60°C for 30 min to mitigate biohazard risk while maintaining RNA quality, then equilibrated to 4°C. Aliquots of 4.875 mL were then deposited into separate wells of a KingFisher 24 deep-well plate (Thermo Fisher). A nuclease-free water negative control was included for the concentration and extraction process. Each sample well was spiked with 5 μL inactivated encapsulated whole vaccine-strain bovine coronavirus (BCoV, Bovilis® Coronavirus vaccine; ∼1.3 × 10^8^ gc/mL by RT-ddPCR), which is a subtype of the species Betacoronavirus 1. All wells received 50 μL Nanotrap® Enhancement Reagent 1 (Ceres Nanosciences, ER1 SKU #10111-30), followed by 75 μL Nanotrap® Magnetic Virus Particles (Ceres Nanosciences, Inc). To concentrate viruses, we applied a Ceres Nanosciences protocol with minor modifications (described in Muralidharan et al. 2025) using a KingFisher Apex robot (Thermo Scientific). RNA and DNA were then extracted from the sample concentrates using the KingFisher Apex according to MagMAX Microbiome Ultra Nucleic Acid Isolation kit (Thermo Fisher) using a modified script to match the input volume and 100 μL elution. Extracts were stored at −80 °C.

### Settled solids sample processing

Settled solids were obtained from each ∼1 L influent sample by allowing solid material to settle in a beaker for at least 30 min and decanting the supernatant. Settled solids were then transferred to centrifuge tubes and centrifuged at 24,000 × g for 30 min at 4 °C for dewatering. Overlying water was carefully decanted. For samples processed in April 2022, four or five aliquots of the resultant dewatered solids (∼75 mg each) were transferred to 50 mL conical tubes and weighed. For samples processed in November 2022, a single ∼750 mg aliquot was used. A stock solution of BCoV (Calf-Guard Cattle Vaccine, PBS Animal Health) was prepared in DNA/RNA Shield (Zymo Research) at a concentration of 500,000 genome copies/mL. This spike stock solution was pipetted into each tube of dewatered solids to a concentration of 1 mL DNA/RNA shield per 75 mg dewatered solids. The spiked dewatered solids were homogenized in conical tubes by bead beating for 2 min at 1,000 rpm using a SPEX™M SamplePrep 2010 Geno/Grinder™ with SPEX 5/32” Grinding Beads and centrifuged for 5 min at 5250 x g. RNA and DNA in the resulting supernatant were extracted using the MagMAX™ Microbiome Ultra Nucleic Acid Isolation Kit (Applied Biosystems by Thermo Fisher Scientific) with use of a KingFisher Flex following manufacturer protocols (Pub. No. MAN0018071 Rev. C.0). Nucleic acid extraction was followed by PCR inhibitor removal using the Zymo™ OneStep-96 PCR Inhibitor Removal Kit (Zymo Research) following manufacturer protocols, with minor modifications (described by Kadonsky et al. 2023). Extracts were stored at −80 °C.

### Library preparation and sequencing

Sample extracts were quantified using the Qubit High Sensitivity RNA and dsDNA kits prior to library preparation. NATtrol™ Influenza A H1 (Flu A) and NATtrol™ Adenovirus Type 1 (Adeno) were purchased from Zepto Metrix (SKU: NATFLUAH1-STQ, NATADV1-STQ) and extracted using the MagMax Ultra Microbiome Kit on a Kingfisher Apex instrument. Input for each sample in library prep was 100 ng total nucleic acids. Controls and Sample extracts received spike-ins at Step 1 of the TurboRNA protocol (3 μL Flu A + 3 μL Adeno + 100 ng sample = 7.5 μL input volume). The libraries were prepared using the provided Beta Illumina protocol^1^, which is designed to be able to use an input sample of mixed DNA/RNA. Because the libraries were to go through enrichment, ribodepletion was not done during library prep. The Turbo RNA library prep method is now called the Illumina Rapid Viral Surveillance Panel v2 Kit.

The libraries prior to enrichment were stored at -80 °C. Quantification was performed using the Qubit dsDNA BR kit, and 200 ng of each library was enriched using the Illumina Virus Surveillance Panel (VSP) v1 according to the provided protocol. Later, the same pre-enrichment library was enriched using VSP v2 by Illumina. Enriched libraries were quantified using the Qubit dsDNA HS kit, and fragment analysis was performed at the UC Davis Genome Center DNA Technologies Core using a Bioanalyzer. Libraries were pooled at equimolar concentrations in two batches (one for VSP v1 and one for VSP v2). The VSP v1 enriched samples were sequenced using the NextSeq 300 cycles PE150 MID output (∼120 million read pairs per lane, ∼6.7 ± 4.7 million per sample) at the UC Davis Genome Center DNA Technologies Core. The VSP v2 enriched set was sequenced using the NovaSeq X PE150 10B (1.2 billion read pairs per lane, ∼66.7 ± 14 million per sample) at the UC Davis Genome Center DNA Technologies Core.

### Computational pipeline

The samples were received as fastq-formatted files (Table S6). Low-quality read removal and adapter trimming were performed using fastp v0.24.0 (Chen 2023) with default parameters. Then, for human decontamination, the reads were mapped to HG19 using Bowtie2 v2.5.4 (Langmead and Salzberg 2012), and the unaligned reads made it to the next stage. Abundances were calculated using Kraken2 (Wood et al. 2019) with the standard database from 09-04-2024, and then with Bracken (Lu et al. 2017) using a minimal read threshold of 10. Post-processing included retaining only the viral species and leaving out sequences associated with positive control spikes from all downstream reporting (Table S1). Reads from the spiked-in positive control viruses did not match the reference genomes with 100% identity. Reads mapped to the most similar representative in the reference dataset, sometimes more than one. Deciding on which species in our datasets were spiked-in was a best-effort decision based on visual inspection of the abundance plots.

Human-targeting viruses were determined using RefSeq’s metadata, selecting all entries with “*Homo sapiens*” in their host field. Matching VSP-enriched viruses with Kraken results was done using a graph data structure (Table S3). Each entry in the two versions of the VSP metadata table added edges between the virus name and its genome accessions. In Kraken’s database, each entry added edges between the species name, its taxon IDs and its genome accessions. PERMANOVA was done using the adonis2 function from the “Vegan” package version 2.6-8 (Oksanen et al. 2025), with 1M permutations and by=”margin”. Rarefaction was done on the raw read counts using the biostuff library v1.1 (Lavon 2025a). Coverage plots were generated using Bowtie2 v2.5.4 to map each sample to each species, using the ‘--very-fast’ mode. The successfully mapped reads were collected, summed up and plotted using an in-house script. Mapping to GreenGenes was performed using database version 2024.09 and Bowtie2 in ‘--very-fast’ mode. Algorithms and data structures were imported from the gostuff library v1.1 (Lavon 2025b). Parsers for biological data files were imported from the biostuff library v1.1.

## Acknowledgements

The wastewater samples were collected and processed within the framework of Healthy Central Valley Together (HCVT), research and public health initiative at the University of California (UC) Davis and Merced focused on increasing access to public health data through wastewater testing in California’s Central Valley. HCVT is committed to health equity, working closely with rural, agricultural, and disadvantaged communities to enable access to critical public health information and resources. We thank HCVT partners and collaborators who contributed to this study.

## Data availability

The sequencing data presented in this study, with adapters and human reads removed, are openly available in the National Center for Biotechnology Information (NCBI) Sequence Read Archive (SRA) under the accession number PRJNA1308571. The code for this project is available on GitHub: github.com/fluhus/sequoia

## Footnotes

1 Illumina RNA Prep with Enrichment, (L) Tagmentation Reference Guide (1000000124435) - Illumina Turbo RNA Prep with Enrichment Reference Guide_BetaV2

