## Supplementary Information for "Performance comparison of probe-capture enrichment sequencing of viruses in wastewater influent and settled solids from California’s Central Valley"

### Supplementary Tables

### Table S1: Positive control spikes and associated species that were removed from presented data where noted.

| Positive Control | Source | Species associated with Positive Control Spikes (removed from presented data where noted) |
| --- | --- | --- |
| Bovine coronavirus (BCoV) (added to each sample before processing) | Betacoronavirus 1 (BCoV, Bovilis® Coronavirus vaccine) | Betacoronavirus 1 |
| Influenza A  (added to the DNA & RNA extract for each sample) | NATtrol™ Influenza A H1 (Flu A), Zepto Metrix (SKU: NATFLUAH1-STQ) | Alphainfluenzavirus influenzae |
| Adenovirus Type 1  (added to the DNA & RNA extract for each sample) | NATtrol™ Adenovirus Type 1 (Adeno), Zepto Metrix (SKU: NATADV1-STQ) | Human mastadenovirus C |

### Table S2: Number of samples in which Positive control spikes viruses were detected.

| Spike Virus | v1 Solid | v1 Influent | v2 Solid | v2 Influent |
| --- | --- | --- | --- | --- |
| Betacoronavirus 1 | 9 | 9 | 9 | 9 |
| Alphainfluenzavirus influenzae | 8 | 8 | 8 | 8 |
| Human mastadenovirus C | 9 | 7 | 9 | 9 |

### Table S3: VSP v2 panel targets detected in study samples. Viral species targeted for enrichment by the VSP v2 panel and detected in at least one sample are listed with the number of samples in which each species was detected (n = 9 per sample group). Entries that were clustered together in the graph analysis are in the same line here. Sequences derived from spiked positive controls were included in the detection counts.

| Virus | v1 Solid | v1 Influent | v2 Solid | v2 Influent |
| --- | --- | --- | --- | --- |
| Adeno-associated virus 2 | 0 | 0 | 9 | 9 |
| Aichi virus 1 | 2 | 4 | 3 | 4 |
| BK polyomavirus | 8 | 9 | 9 | 9 |
| Coxsackievirus B (species Enterovirus B) | 0 | 0 | 0 | 1 |
| Enterovirus A | 0 | 4 | 4 | 7 |
| Enterovirus C | 0 | 0 | 1 | 2 |
| Enterovirus D | 0 | 0 | 0 | 2 |
| Epstein-Barr virus | 0 | 0 | 7 | 5 |
| Herpes simplex virus 1 | 0 | 0 | 5 | 3 |
| Human adenovirus A | 3 | 1 | 9 | 8 |
| Human adenovirus B | 2 | 0 | 9 | 6 |
| Human adenovirus C | 9 | 7 | 9 | 9 |
| Human adenovirus D | 5 | 1 | 9 | 8 |
| Human adenovirus F | 9 | 7 | 9 | 9 |
| Human adenovirus G | 1 | 0 | 9 | 7 |
| Human coronavirus 229E | 1 | 4 | 1 | 3 |
| Human coronavirus HKU1 | 0 | 8 | 0 | 6 |
| Human coronavirus NL63 | 0 | 0 | 0 | 3 |
| Human coronavirus OC43 | 9 | 9 | 9 | 9 |
| Human cytomegalovirus | 0 | 0 | 3 | 0 |
| Human papillomavirus 16 | 1 | 0 | 1 | 1 |
| Human papillomavirus 53 | 1 | 0 | 0 | 1 |
| Human papillomavirus 6 | 0 | 0 | 1 | 0 |
| Human parainfluenza virus 1 | 0 | 3 | 0 | 2 |
| Human parainfluenza virus 2 | 0 | 1 | 0 | 1 |
| Human parainfluenza virus 3 | 1 | 6 | 0 | 6 |
| Human parainfluenza virus 4 | 0 | 2 | 0 | 0 |
| Human parechovirus | 0 | 0 | 0 | 1 |
| Human polyomavirus 6 | 2 | 8 | 2 | 8 |
| Human polyomavirus 7 | 0 | 3 | 0 | 3 |
| Human respiratory syncytial virus A | 0 | 4 | 1 | 3 |
| Influenza A virus | 8 | 8 | 8 | 8 |
| Influenza C virus | 0 | 0 | 0 | 2 |
| JC polyomavirus | 9 | 9 | 9 | 9 |
| LI polyomavirus | 0 | 0 | 1 | 2 |
| MW polyomavirus | 0 | 0 | 0 | 4 |
| Mamastrovirus 1 | 8 | 8 | 9 | 9 |
| Mamastrovirus 6 | 0 | 0 | 8 | 9 |
| Mamastrovirus 8 | 0 | 0 | 7 | 8 |
| Mamastrovirus 9 | 0 | 0 | 5 | 4 |
| Merkel cell polyomavirus | 9 | 9 | 9 | 9 |
| Norovirus, Norovirus GI, Norovirus GII, Norovirus GIV | 8 | 9 | 9 | 9 |
| Rotavirus A | 3 | 0 | 6 | 2 |
| Salivirus A | 0 | 0 | 3 | 3 |
| Sandfly fever Sicilian virus | 0 | 0 | 1 | 0 |
| Sapovirus | 2 | 8 | 9 | 9 |
| Severe acute respiratory syndrome coronavirus 2, Severe acute respiratory syndrome coronavirus | 6 | 9 | 6 | 9 |
| Torque teno virus | 0 | 0 | 0 | 2 |
| Trichodysplasia spinulosa-associated polyomavirus | 0 | 2 | 0 | 2 |
| Varicella-zoster virus | 0 | 0 | 1 | 0 |
| WU polyomavirus | 3 | 5 | 3 | 7 |

### Table S4: Genome types of phyla for identified viruses (used for calculating DNA/RNA content), annotated manually. The 20 most abundant viruses are listed next to their phyla.

| Phylum | Genome Type | Top 20 most abundant species |
| --- | --- | --- |
| Cossaviricota | DNA | Betapolyomavirus secuhominis, Betapolyomavirus hominis, Dependoparvovirus primate1, Alphapolyomavirus quintihominis |
| Cressdnaviricota | DNA |  |
| Duplornaviricota | RNA |  |
| Hofneiviricota | DNA |  |
| Kitrinoviricota | RNA |  |
| Lenarviricota | RNA |  |
| Negarnaviricota | RNA | Alphainfluenzavirus influenzae |
| Nucleocytoviricota | DNA |  |
| Peploviricota | DNA |  |
| Phixviricota | DNA |  |
| Pisuviricota | RNA | Betacoronavirus 1, Human astrovirus, Norwalk virus, Mamastrovirus 1, Rabbit coronavirus HKU14 |
| Preplasmiviricota | DNA | Human mastadenovirus C, Human mastadenovirus F, Simian mastadenovirus F, Human mastadenovirus D, Simian mastadenovirus E, Human mastadenovirus A  Uroviricota: Jouyvirus ev017, Carjivirus communis, Koutsourovirus KDA1, Carjivirus hominis |
| Uroviricota | DNA |  |

### Table S5: VSP v2 viruses detected. Virus names from the VSP v2 documentation, that were found in our data by name search.

| Virus Name | Genome Type |
| --- | --- |
| Adeno-associated virus 2 | DNA |
| Aichi virus 1 | RNA |
| BK polyomavirus | DNA |
| Enterovirus A | RNA |
| Enterovirus B | RNA |
| Enterovirus D | RNA |
| Epstein-Barr virus | DNA |
| Herpes simplex virus 1 | DNA |
| Human adenovirus A | DNA |
| Human adenovirus B | DNA |
| Human adenovirus C | DNA |
| Human adenovirus D | DNA |
| Human adenovirus F | DNA |
| Human adenovirus G | DNA |
| Human coronavirus 229E | RNA |
| Human coronavirus HKU1 | RNA |
| Human coronavirus NL63 | RNA |
| Human coronavirus OC43 | RNA |
| Human cytomegalovirus | DNA |
| Human papillomavirus 16 | DNA |
| Human papillomavirus 53 | DNA |
| Human papillomavirus 6 | DNA |
| Human parainfluenza virus 1 | RNA |
| Human parainfluenza virus 2 | RNA |
| Human parainfluenza virus 3 | RNA |
| Human parainfluenza virus 4 | RNA |
| Human parechovirus | RNA |
| Human polyomavirus 6 | DNA |
| Human polyomavirus 7 | DNA |
| Human respiratory syncytial virus A | RNA |
| Influenza A virus | RNA |
| Influenza C virus | RNA |
| JC polyomavirus | DNA |
| LI polyomavirus | DNA |
| MW polyomavirus | DNA |
| Mamastrovirus 1 | RNA |
| Mamastrovirus 6 | RNA |
| Mamastrovirus 8 | RNA |
| Mamastrovirus 9 | RNA |
| Merkel cell polyomavirus | DNA |
| Norovirus | RNA |
| Poliovirus (Enterovirus Species C) | RNA |
| Rotavirus A | RNA |
| Salivirus A | RNA |
| Sandfly fever Sicilian virus | RNA |
| Sapovirus | RNA |
| Severe acute respiratory syndrome coronavirus 2 | RNA |
| Torque teno virus | DNA |
| Trichodysplasia spinulosa-associated polyomavirus | DNA |
| Varicella-zoster virus | DNA |
| WU polyomavirus | DNA |

### Table S6. Summary of sample sequencing quality. Three timepoints from 3 WWTPs were assessed with liquid wastewater (9 samples) and solids (9 samples), using both VSP v1 and v2 enrichment approaches, yielding 36 sequenced samples.

| Sample ID | Collection Date | Panel Version | Type | Location | Number of reads (raw) | Number of reads (QC) | %QC |
| --- | --- | --- | --- | --- | --- | --- | --- |
| A1 | 4/14/2022 | 1 | Solid | Site A | 4238964 | 4025997 | 94.98% |
| A2 | 11/16/2022 | 1 | Solid | Site A | 6680641 | 6357805 | 95.17% |
| A3 | 11/30/2022 | 1 | Solid | Site A | 4734475 | 4534663 | 95.78% |
| B1 | 4/7/2022 | 1 | Solid | Site B | 4773720 | 4549266 | 95.30% |
| B2 | 11/16/2022 | 1 | Solid | Site B | 9986499 | 9464167 | 94.77% |
| B3 | 11/30/2022 | 1 | Solid | Site B | 9044103 | 8566549 | 94.72% |
| C1 | 4/15/2022 | 1 | Solid | Site C | 3139635 | 2993833 | 95.36% |
| C2 | 11/16/2022 | 1 | Solid | Site C | 3227548 | 3008581 | 93.22% |
| C3 | 11/30/2022 | 1 | Solid | Site C | 6292457 | 6048018 | 96.12% |
| A1 | 4/14/2022 | 1 | Influent | Site A | 8732614 | 8367161 | 95.82% |
| A2 | 11/16/2022 | 1 | Influent | Site A | 7445184 | 7100594 | 95.37% |
| A3 | 11/30/2022 | 1 | Influent | Site A | 10279575 | 9828680 | 95.61% |
| B1 | 4/7/2022 | 1 | Influent | Site B | 15355572 | 14783177 | 96.27% |
| B2 | 11/16/2022 | 1 | Influent | Site B | 8767585 | 8373859 | 95.51% |
| B3 | 11/30/2022 | 1 | Influent | Site B | 18310127 | 17597000 | 96.11% |
| C1 | 4/15/2022 | 1 | Influent | Site C | 1783262 | 1731504 | 97.10% |
| C2 | 11/16/2022 | 1 | Influent | Site C | 15557176 | 14927609 | 95.95% |
| C3 | 11/30/2022 | 1 | Influent | Site C | 13787275 | 13342339 | 96.77% |
| A1 | 4/14/2022 | 2 | Solid | Site A | 64273188 | 63702945 | 99.11% |
| A2 | 11/16/2022 | 2 | Solid | Site A | 85200656 | 84396090 | 99.06% |
| A3 | 11/30/2022 | 2 | Solid | Site A | 64683482 | 64015977 | 98.97% |
| B1 | 4/7/2022 | 2 | Solid | Site B | 59664782 | 58683504 | 98.36% |
| B2 | 11/16/2022 | 2 | Solid | Site B | 62387162 | 61353320 | 98.34% |
| B3 | 11/30/2022 | 2 | Solid | Site B | 83638393 | 82569444 | 98.72% |
| C1 | 4/15/2022 | 2 | Solid | Site C | 51657894 | 50863969 | 98.46% |
| C2 | 11/16/2022 | 2 | Solid | Site C | 58590390 | 57864075 | 98.76% |
| C3 | 11/30/2022 | 2 | Solid | Site C | 80386810 | 79563619 | 98.98% |
| A1 | 4/14/2022 | 2 | Influent | Site A | 69454573 | 68764536 | 99.01% |
| A2 | 11/16/2022 | 2 | Influent | Site A | 89295618 | 88513412 | 99.12% |
| A3 | 11/30/2022 | 2 | Influent | Site A | 94870765 | 94020182 | 99.10% |
| B1 | 4/7/2022 | 2 | Influent | Site B | 82380601 | 81245245 | 98.62% |
| B2 | 11/16/2022 | 2 | Influent | Site B | 79749189 | 78790903 | 98.80% |
| B3 | 11/30/2022 | 2 | Influent | Site B | 65052639 | 64569834 | 99.26% |
| C1 | 4/15/2022 | 2 | Influent | Site C | 84714702 | 83856220 | 98.99% |
| C2 | 11/16/2022 | 2 | Influent | Site C | 97058549 | 96091402 | 99.00% |
| C3 | 11/30/2022 | 2 | Influent | Site C | 91008373 | 90188036 | 99.10% |

### Supplementary Figures

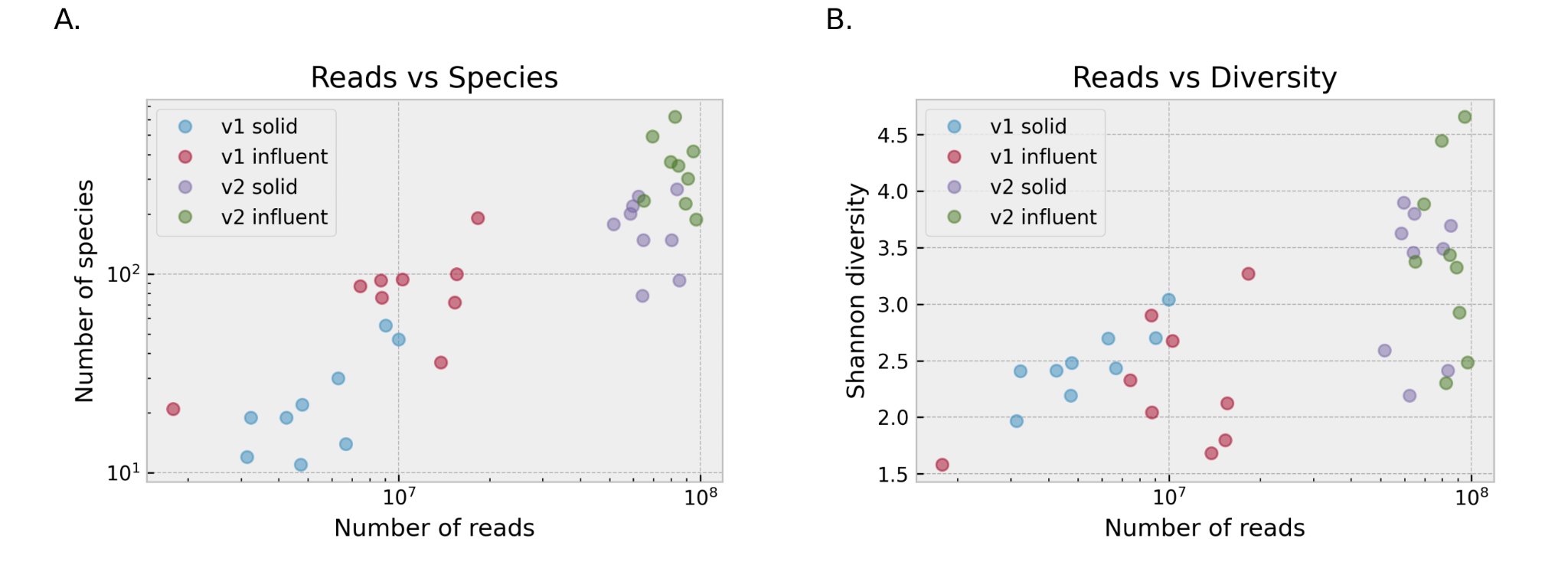

***Figure S1:*** *Number of total reads (including spike) compared with (A) number of viral species identified by Kraken; and with (B) viral Shannon diversity (post spike removal).*

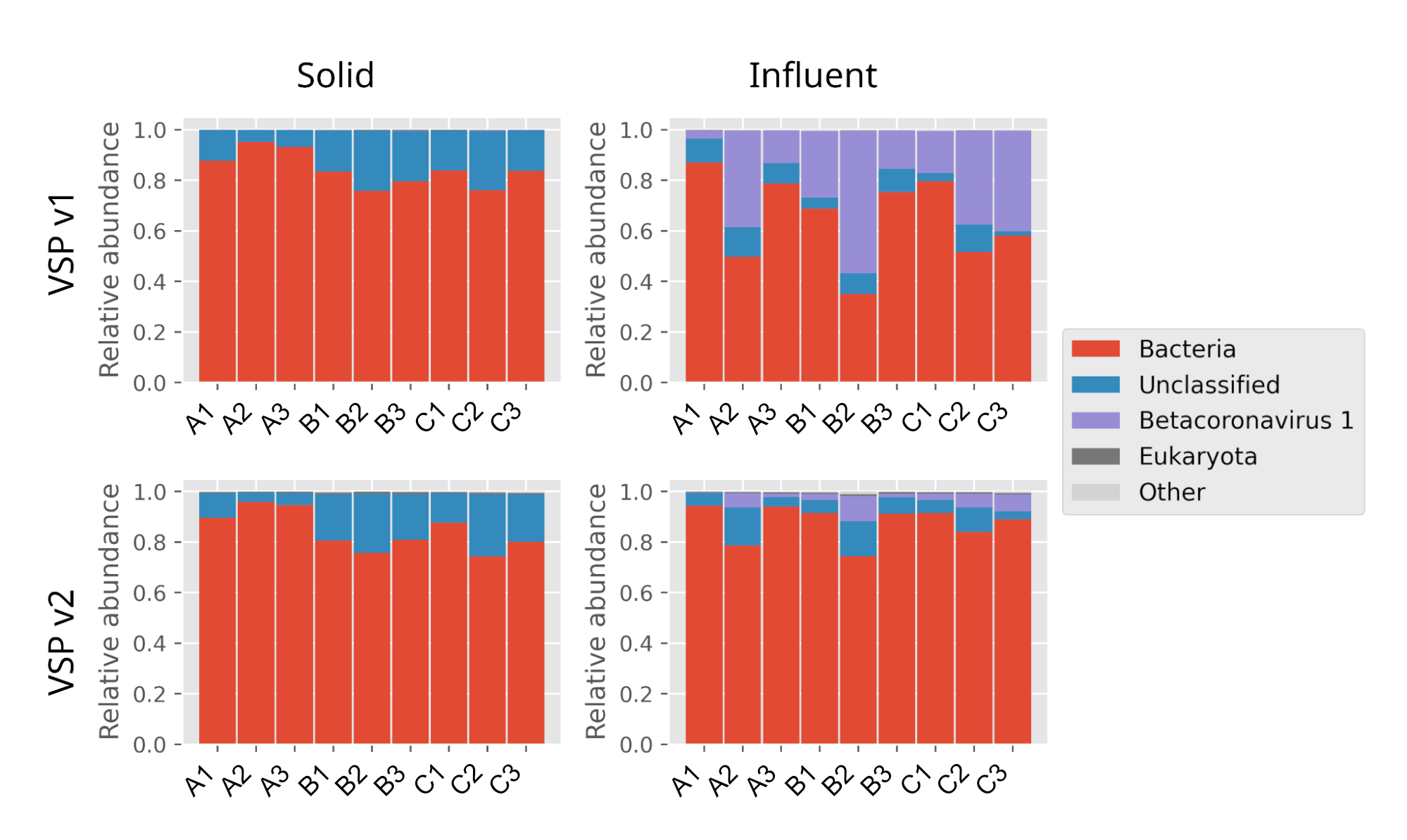

***Figure S2:*** *Raw read assignment distributions among major taxonomic assignment groups, including spikes and non-viral groups. Betacoronavirus 1 represents the bovine coronavirus process-control spike. Viruses account for 8 ± 14% reads per sample.*

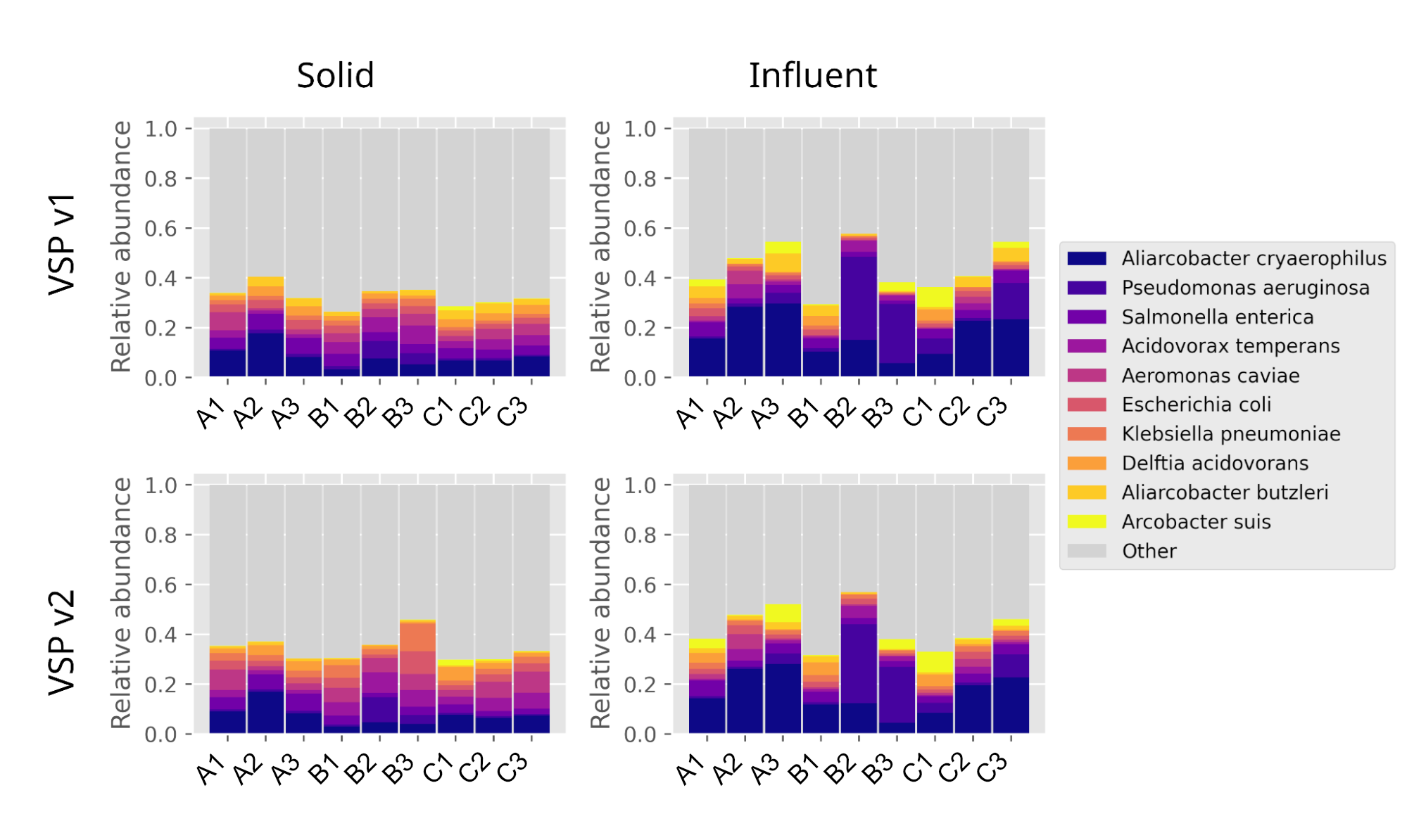

***Figure S3:*** *Bacterial relative abundances in samples. The top 10 species by average relative abundance are colored.*

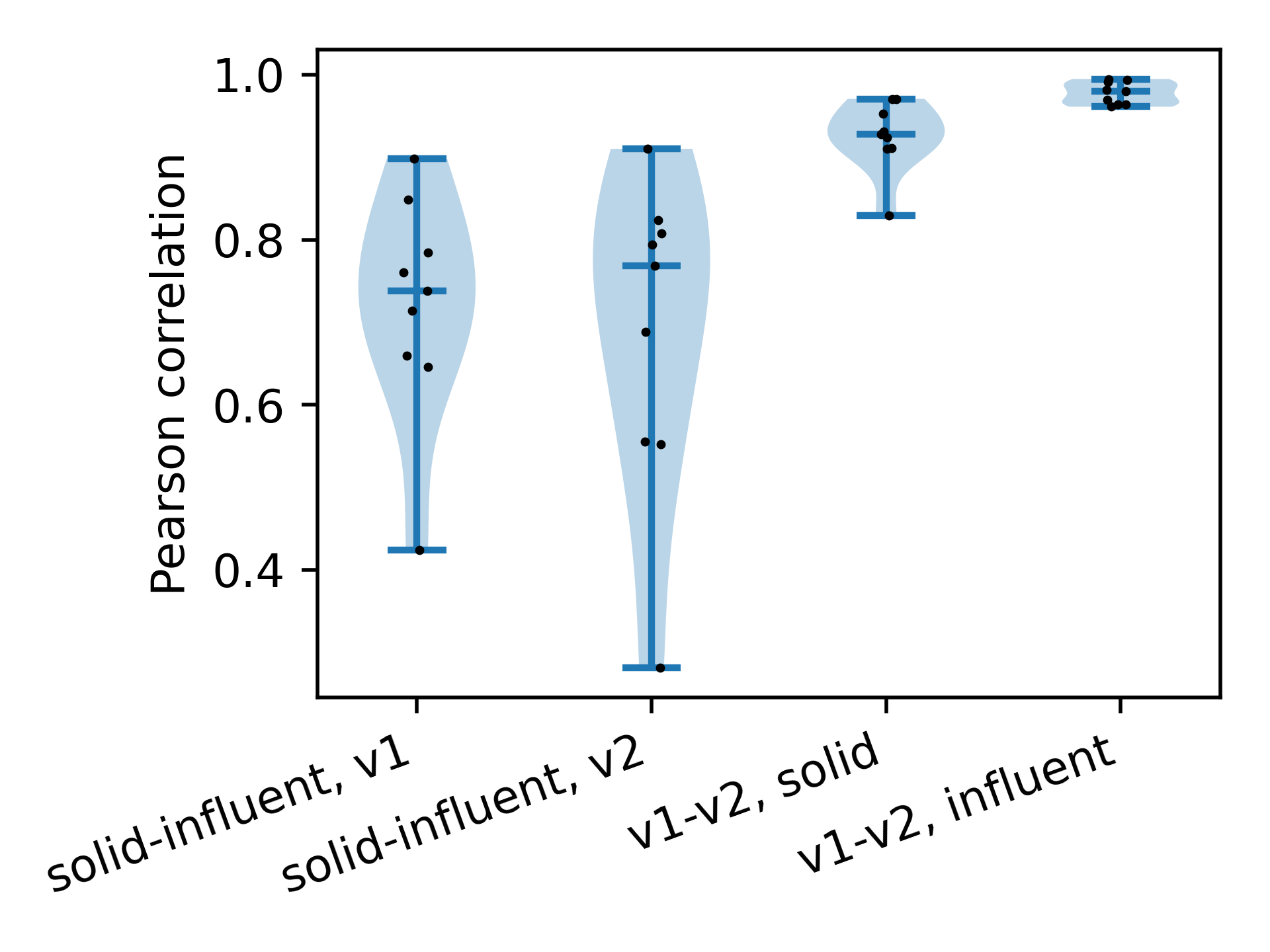

***Figure S4:*** *Bacterial relative abundance correlations across sample types. Each point represents a single sample, comparing two sequencing methods.*

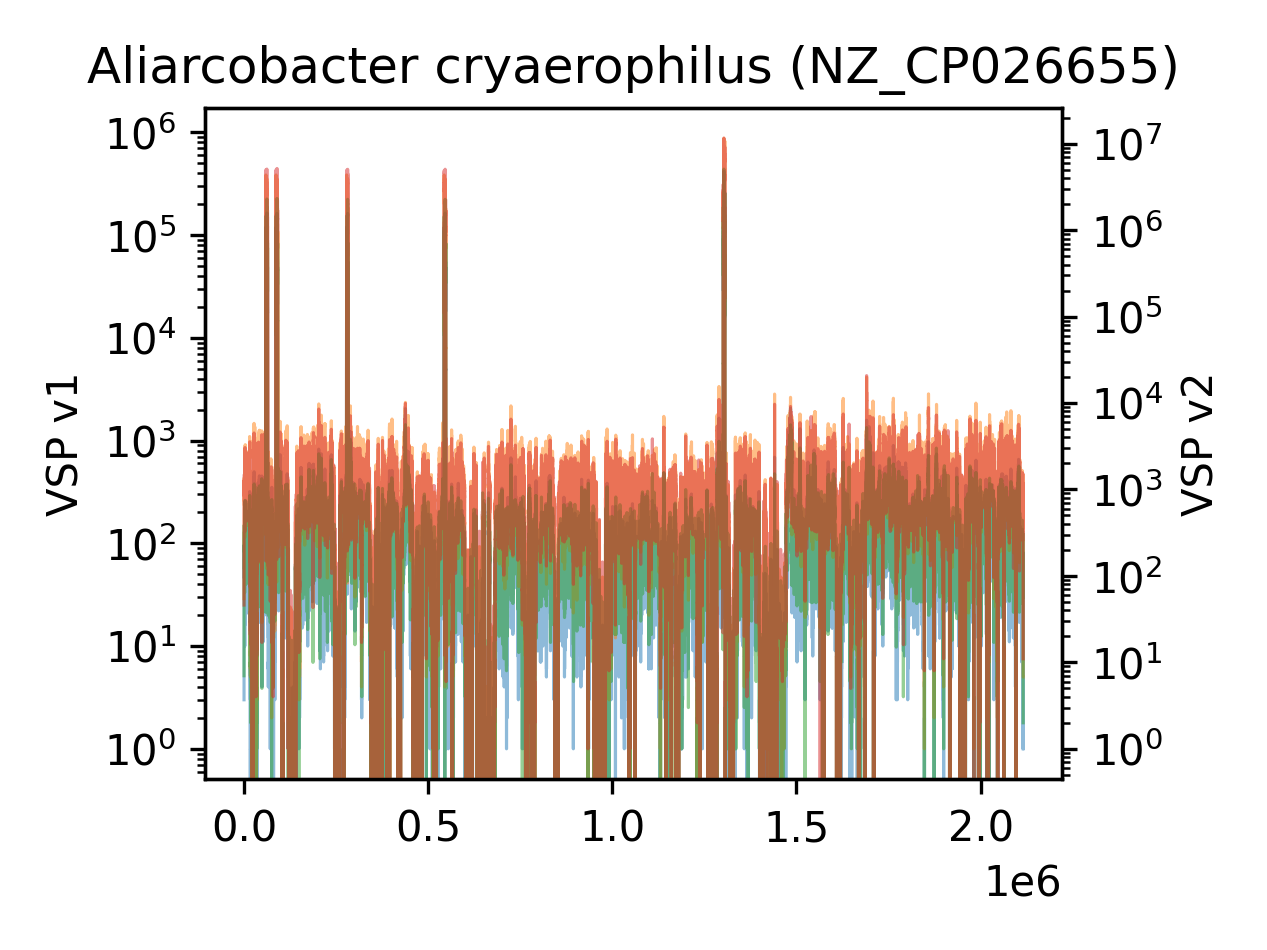

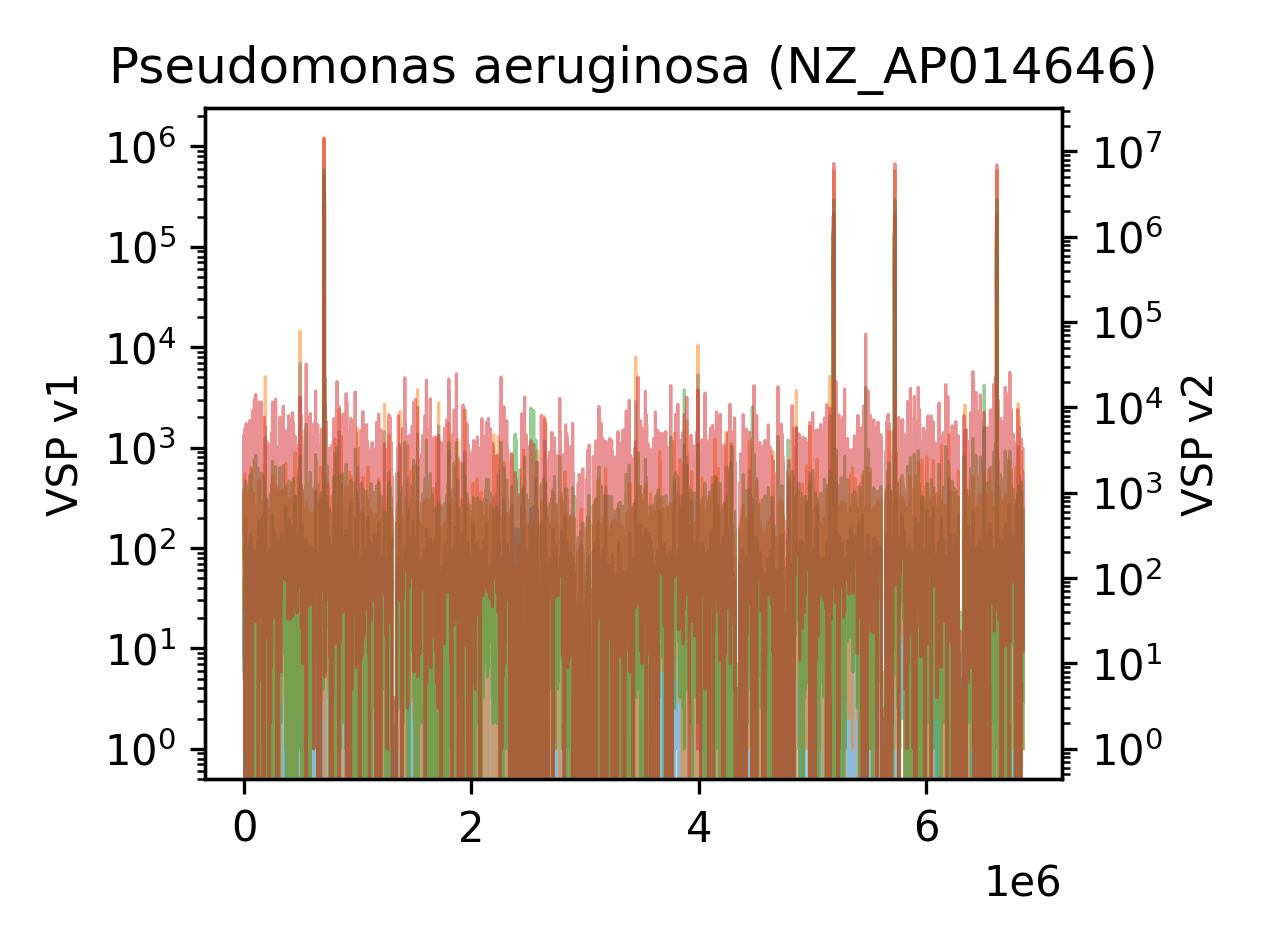

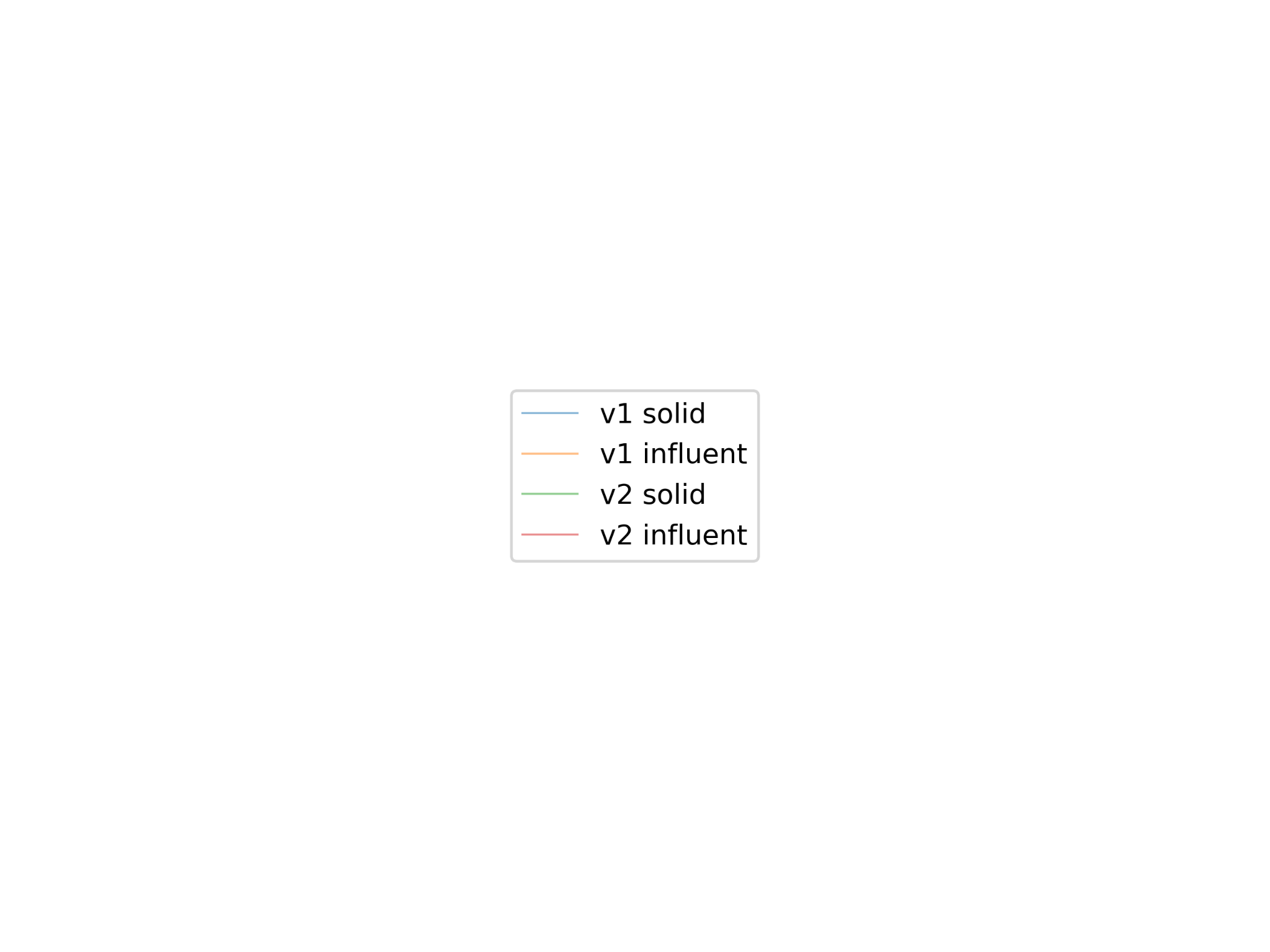

***Figure S5:*** *Genome coverage of the two most abundant bacteria, genome accession code in parentheses. All reads from all samples were mapped to each genome, and grouped by sample type. The Y axes are log-scaled.*
